# Loss of dystrophin reduces CB1 receptor expression and endocannabinoid-dependent synaptic plasticity in the cerebellar cortex

**DOI:** 10.64898/2026.03.20.713279

**Authors:** Emily Averyt, Shaarang Mitra, Jason R. Pugh

## Abstract

Duchenne Muscular Dystrophy (DMD) is a debilitating degenerative condition with complex musculoskeletal and cognitive symptoms. The protein responsible, dystrophin, is expressed in both muscle tissue and within the central nervous system (CNS) where it localizes to inhibitory synapses. Recent work has shown that dystrophin loss in skeletal muscle leads to abnormalities in endocannabinoid signaling, particularly related to Cannabinoid Receptor Type 1 (CB1R) signaling pathways. CB1Rs are highly expressed throughout the CNS, and have been implicated in short- and long-term plasticity mechanisms. Despite this curious overlap, no work examines how dystrophin loss impacts CB1R signaling in the CNS, a mechanism that may contribute to the diverse neurological pathologies seen in DMD patients. To address this, we used a combination of immunofluorescent labeling and ex vivo electrophysiology to examine CB1R signaling at three classes of synapses within the cerebellum. Utilizing DMD^mdx^ mice, a mouse model of DMD, we find that loss of dystrophin significantly impairs CB1R signaling specifically at parallel fiber-Purkinje Cell synapses, a key location for cerebellar learning. We also find that endocannabinoid-mediated long-term depression at these synapses is absent. Loss of endocannabinoid signaling and synaptic plasticity may contribute to cerebellar dysfunction and motor control symptoms in DMD. These data suggest that dystrophin loss may have previously undescribed consequences for CNS function, and that modulation of endocannabinoid signaling may be a therapeutic strategy for symptom management.

**Significance Statement:** Duchenne Muscular Dystrophy (DMD) is a degenerative condition with severe CNS deficits in addition to the well-known muscle weakening. However, no effective treatments currently exist for CNS-related aspects of this disease. Given that endocannabinoid signaling is altered in dystrophic muscle and the importance of endocannabinoid signaling in CNS function, we examined endocannabinoid signaling in the cerebellum of DMD^mdx^ mice, a model of DMD. Utilizing immunolabeling and ex vivo electrophysiology, we find a significant decrease in CB1R expression and functionality specifically at parallel fiber synapses, resulting in reduced or abolished short- and long-term synaptic plasticity. These findings demonstrate that changes in endocannabinoid function contribute to CNS deficits in DMD and open the door to new potential therapeutic targets for treatment.

## INTRODUCTION

Duchenne muscular dystrophy (DMD) is a severe, degenerative musculoskeletal disease that impacts 1 in 5000 males. The condition is caused by mutations in the X-linked *DMD* gene, leading to loss of dystrophin protein. Cases are diagnosed with early coordination and postural issues, developing into progressive muscular degeneration. In addition to muscular symptoms, DMD patients exhibit severe, non-progressive cognitive deficits, suggesting the disease may also impact central nervous system (CNS) function (Duchenne, 1868). DMD patients are commonly reported to have reduced working memory (Hinton et al., 2000), intellectual disability (average full-scale IQ ∼80) (Billard et al., 1992; Cotton et al., 2005; Ricotti et al., 2016; Doorenweerd et al., 2017), and impaired verbal expression (Karagan et al., 1980). DMD patients also show increased rates of neurodevelopmental disorders including autism-spectrum disorders, attention-deficit/hyperactivity disorder, and obsessive-compulsive disorder (Vaillend et al., 2025). These factors point to a significant understudied impairment in CNS function and development.

In the CNS, full length (dp427) dystrophin is found at specific inhibitory synapses within various brain regions, including the cerebellum (Lidov et al., 1993), where it links the transynaptic dystrophin-glycoprotein complex to the actin cytoskeleton. Dystrophin is highly expressed in cerebellar Purkinje Cells (PCs), where it localizes to the postsynaptic densities of inhibitory synapses between molecular layer interneurons (MLIs) and PCs (Knuesel et al., 1999). PCs from DMD^mdx^ mice (a model of DMD lacking dystrophin expression) display a significant reduction in the number, size, and strength of inhibitory synapses. Taken together, these deficits reduce synaptic input into PCs by ∼50%, and result in impaired motor learning (Wu et al., 2022; Prigogine et al., 2024).

In addition to deficiencies in muscle function and GABAergic signaling, DMD may also impact endocannabinoid signaling. Recent studies found alterations in Cannabinoid Receptor Type 1 (CB1R) expression in muscle stem cells, which may impact regenerative capabilities and exacerbate disease progression (Iannotti et al., 2018, 2019). CB1Rs are also one of the most abundant GPCRs in the brain, where they modulate a range of circuits and plasticity mechanisms (Kendall and Yudowski, 2017; Winters and Vaughan, 2021). CB1Rs are expressed at the presynaptic terminals of both excitatory and inhibitory synapses, where they primarily inhibit voltage-gated calcium channels and reduce transmitter release. In addition to this short-term plasticity, CB1R activation is also necessary for various forms of long-term plasticity (Winters and Vaughan, 2021). In the cerebellar circuit, CB1R activity is necessary for LTD/LTP induction at Parallel Fiber (PF) to Purkinje Cell (PC) synapses (Safo and Regehr, 2005; Carey et al., 2011; Wang et al., 2014), a critical site of plasticity for motor learning, coordination, and cognitive aspects of cerebellar function (Lisberger, 1998). Based on observed changes in peripheral CB1Rs, the high expression of CB1Rs and dystrophin in the cerebellar cortex, and the role of CB1Rs in regulating synaptic plasticity, we hypothesize that changes in CB1R expression may contribute to cerebellar dysfunction and the prevalence of neurodevelopmental disorders in DMD patients.

To test this hypothesis we examined expression patterns and functional alterations in endocannabinoid signaling within the cerebellum utilizing the DMD^mdx^ mouse model. Through immunofluorescent labeling and ex vivo electrophysiology, we examine three classes of synapses within the cerebellum reported to express CB1Rs: Molecular Layer Interneuron (MLI)-PC, PF-PC, and Climbing Fiber (CF)-PC synapses. Despite dystrophin expression being limited to MLI-PC synapses, we did not find changes in CB1R expression or CB1R-dependent short-term plasticity at these synapses. Instead, CB1R expression, measured by immunofluorescence, CB1R-dependent short-term plasticity, and response to a CB1R agonist, was significantly reduced at PF-PC synapses. CB1R expression at CF synapses was modest, with little or no change across genotype. Finally, we show that reduced CB1R expression at PF-PC synapses is associated with loss of LTD expression, suggesting plasticity dynamics are heavily altered in the cerebellar circuit. Together, these data point to significant alterations in endocannabinoid signaling as a result of impaired dystrophin expression within the DMD^mdx^ mouse model, which may impact broader cerebellar functions.

## METHODS

### Animals

All experimental procedures involving animals were approved by the Institutional Animal Care and Use Committee at UT Health San Antonio. Electrophysiology and immunofluorescent labeling experiments utilized male DMD^mdx^ mice (C57BL/10ScSn-Dmd*^mdx^*/J; JAX #001801) or WT littermate controls. Immunofluorescent labeling experiments also included CB1-KO animals (cnr1−/−, NIMH transgenic core, NIH, Bethesda, MD) provided by Chu Chen at University of Texas Health Science Center at San Antonio. Only male mice were used to better replicate the etiology of Duchenne Muscular Dystrophy, an X-linked recessive disorder primarily impacting male patients. Animals were kept on a 12 hr light/dark cycle with *ad libitum* access to food and water. Investigators were blinded to genotype during data acquisition and analysis.

### Ex vivo patch-clamp electrophysiology

#### Slice preparation

Acute parasagittal brain slices were prepared from P30-50 WT and DMD^mdx^ mice the day of each recording session. Mice were deeply anesthetized with isoflurane inhalation before decapitation and rapid dissection of cerebellar tissue. The cerebellum was placed in ice-cold oxygenated artificial cerebrospinal fluid (ACSF) containing the following (in mM): 119 NaCl, 26.2 NaHCO_3_, 2.5 KCl, 1.0 NaH_2_PO_4_, 11 glucose, 2 CaCl_2_, 1.3 MgCl_2_. Parasagittal slices (300 µm) were cut from the vermis of the cerebellum using a vibratome (model VT1200S, Leica Biosystems, IL), then incubated at 34°C for 30 min in oxygenated ACSF. Following recovery, slices were maintained at room temperature for up to 6 hrs prior to use in electrophysiological experiments.

#### Electrophysiological recording

Slices were gently transferred by wide mouth pasteur pipette to a recording chamber perfused with 31-33°C ACSF (flow rate of ∼2 ml/min) and housed in a SliceScope Pro upright microscope (Scientifica Instruments, UK). Voltage clamp recordings were collected from PCs using one of the following internal solutions (in mM). CsCl internal (IPSC recording experiments): 140 CsCl, 10 HEPES, 2 MgCl_2_, 0.16 CaCl_2_, 0.5 EGTA, 4 Na-ATP, 0.5 Na-GTP, 2 QX-314 Bromide (Tocris Bioscience). CsMeSO_3_ internal (EPSC recording experiments): 136 CsMeSO_3_, 10 CsCl, 10 HEPES, 0.2 EGTA, 2 QX-314 Bromide, 4 Na-ATP, 0.3 Na-GTP, and 0.16 CaCl_2_. K-Gluconate internal (PF-LTD experiments): 137 K-Gluconate, 2 KCl, 4 MgCl2, 10 HEPES, 5 EGTA, 4 Na-ATP, 0.5 Na-GTP. All internal solutions were adjusted to pH 7.3-7.4 using KOH or CsOH and osmolarity was adjusted to 285-295 mOsm. Cells were patched using 1.5-4 MΩs borosilicate glass pipettes (Sutter Instrument, CA) that were pulled on a Sutter pipette puller (Model P-100, Sutter Instrument, CA). Electrophysiological currents were recorded with a Multiclamp 700B amplifier (Molecular Devices, CA), filtered at 5 kHz and digitized at 50 kHz (Digidata 1550B, Molecular Devices, CA). Data were collected using pCLAMP software (Molecular Devices, CA). Access resistances of all cells were monitored, and cells deviating from 20±10 MΩs or shifting by >10% were not included..

Whole-cell patch clamp recordings were made from PCs in folia 5 and 6 of the vermis and a glass stimulating electrode containing ACSF was placed 75-150 µm from the patched PC soma in the molecular layer (MLI IPSC and PF EPSC recordings) or near the base of the patched PC (CF EPSC recordings). Synaptic currents were evoked by electrical stimulation through a constant voltage isolated stimulator (∼5-50 V; Digitimer) triggered from pCLAMP software. Stimulus intensity was adjusted to produce IPSC and PF-EPSC amplitudes of 200-400 pA. Paired-pulse ratio (PPR) was measured by delivering pairs of electrical stimulation with a 20 ms ISI. Stimulus response curves were generated by recording EPSCs at the minimum stimulation necessary to generate a ∼200 pA EPSC, followed by recording EPSCs at a stimulus intensity 20%, 60%, 100%, 140%, and 180% of that value.

Where indicated, excitatory post-synaptic currents (EPSCs) were isolated by inclusion of 100 µm picrotoxin (PTX; Abcam Biochemicals) and inhibitory post-synaptic currents (IPSCs) were isolated by inclusion of 5 µm R-CPP (Abcam Biochemicals) and 10 µm NBQX (Thermo Scientific Chemicals) in the bath solution. Additionally, 1 µm WIN-55,212-2 (Enzo Biochem) was used to activate CB1Rs in relevant experiments.

#### Induction of Depolarization-Induced Suppression of Inhibition (DSI) and Excitation (DSE)

For MLI-DSI and PF-DSE experiments, PCs were clamped to −70 mV, and 5 synaptic responses were evoked at 0.5 Hz to establish a baseline amplitude. Synaptic stimulation was paused for 6 seconds and the PC was depolarized to 0 mV for 2 seconds. 0.5 Hz stimulation then resumed for 61 seconds, after which cells were allowed to rest until the next sweep (2.5 minute intersweep interval). For CF-DSE, 0.3 µm NBQX was included in the bath solution (in addition to 100 µm PTX) to partially antagonize AMPA receptors, reducing the amplitude of the CF-EPSC to allow for better voltage-clamp. CF responses were identified by the large amplitude (>1000 pA), extensive paired-pulse depression, and all-or-nothing responses. The stimulus strength was adjusted to the minimum stimulus amplitude required to consistently yield a CF-response. The CF-DSE protocol was similar to that described above, however the stimulus frequency was reduced to 0.25 Hz to minimize short-term depression and vesicle depletion, the PC was depolarization was extended to 4 seconds to maximize 2-AG production/release, and the intersweep interval was extended to 4 minutes to allow recovery between sweeps. 3-6 sweeps were recorded from each cell. DSI/DSE were measured by comparing the amplitude of averaged baseline responses against the amplitude of the first 3 responses after the depolarization phase.

#### WIN 55,212-2 experiments

For experiments testing the effects of WIN 55,212-2 (WIN) on PF or CF EPSCs, PCs were patched and stimulating electrodes were placed as described above. A stable baseline of EPSC amplitudes was obtained by stimulating EPSCs for 3-5 minutes (every 5 or 15 seconds for PF and CF, respectively). Cells that did not show stable EPSC amplitudes during the baseline period were discarded. Following this baseline period, 1 µM WIN, a CB1R agonist, was washed onto the slice by bath perfusion. EPSC stimulation continued through drug wash-on until the responses stabilized again at a lower value (usually 20-25 minutes after WIN application). EPSC amplitude was calculated as the average of 3 minutes of responses once a plateau phase had been reached. Slices were discarded and tubing was rinsed for 30 minutes between recordings.

#### PC LTD induction

A K-Gluconate internal solution (see above) was used for all PC LTD experiments, and 100 µm PTX was included in the bath to prevent GABA_A_R-mediated currents. To evoke PF EPSCs, a stimulating electrode was placed 50-150 µm into the molecular layer, and the stimulus intensity was adjusted to generate 200-400 pA PF responses. A second stimulating electrode was placed 50-100 µm into the granule cell layer, and adjusted to produce a CF response. PCs were voltage clamped to −70 mV, and PFs were stimulated with a 10 Hz paired-pulse protocol (5 seconds intersweep interval) for at least 5 minutes to establish a baseline response amplitude. Following the baseline period, the LTD induction protocol, consisting of 10 PF stimuli at 100 Hz followed immediately by 2 CF stimuli at 20 Hz (Carey et al., 2011), was delivered while holding the cell in current-clamp mode. The LTD induction protocol was repeated every 10 seconds for 5 minutes. Following induction, PCs were returned to voltage-clamp mode and PF stimulation resumed for at least 25 minutes to assess LTD. The magnitude of LTD was calculated by comparing the last five minutes of EPSC recording to the baseline amplitude.

### Immunofluorescent labeling

P50-P60 male mice (4 DMD^mdx^, 4 WT littermates, 1 CNR1^−/−^) were anesthetized with inhaled isoflurane followed by deep anesthesia with Avertin (450 mg/kg, injected IP). Transcardial perfusions were performed using ∼10 mL PBS, followed by 2x bodyweight equivalent volume of 3% glyoxal (Jahncke et al., 2025). The cerebellum was isolated from the skull, submerged in 3% glyoxal solution for 24 hours, then transferred to 30% sucrose solution until saturated (2-3 days). Tissue was subsequently frozen in O.C.T. gel, sliced at 30 µm using a Leica CM 1850 UV Cryostat and affixed to charged glass microscope slides. Slides were stored at −80℃ until staining.

After thawing, slides were washed (PBS, 3x 10 min) and incubated with blocking solution (2% bovine serum albumin, 5% goat serum, 0.3% Triton-X in PBS, 2 hours). Slides were incubated for 48 hours at 4°C under agitation with the following primary antibodies in blocking buffer: calbindin (1:3000, Sigma-Aldrich, Cat# C9848), CB1R (1:1000, Cayman Chemical Company, Cat# 10006590), and one of the following vesicular transporters - VGAT (1:250, Synaptic Systems, Cat# 131004), VGLUT2 (1:500, Synaptic Systems, Cat# 135418), or VGLUT1 (1:5000, Synaptic Systems, Cat# 135304). After primary incubation, slides were once again washed with PBS 3×10 min. Slides were incubated for 2 hours at room temperature under agitation with the following secondary antibodies in blocking buffer: Alexa Fluor 488 goat 𝛼 mouse (1:500, Thermo Fisher, Cat# A11029), Alexa Fluor 568 goat 𝛼 guinea pig (1:200, Thermo Fisher, Cat# A11075), and Alexa Fluor 647 goat 𝛼 rabbit (1:500, Thermo Fisher, Cat# A21245). After secondary incubation, slides were washed a final time with PBS 3×10 min, dried, then covered in SlowFade Gold Antifade Mountant (Invitrogen, Cat# S36937) and sealed with a glass cover slip. Imaging was performed using a Stellaris 8 Confocal Microscope (Leica) at 63x magnification. Z stacks were obtained with a pitch of 0.4 µm. Images were postprocessed using the LIGHTNING Algorithm of the Leica Application Suite X software to reduce noise and out of plane light scattering.

#### Machine Learning Regional identification

A small cohort of 42 images were manually masked for anatomical regions - Molecular Layer, Granule Cell Layer, and Purkinje Cell Layer - based solely on calbindin staining, which is restricted to PC somas and dendrites. A 2D U-Net based convolutional neural network was trained on the calbindin channel of the source image and the provided binary masks. The system used a 70/15/15 train/validation/test split paradigm, resulting in 30 training images, 6 validation, and 6 testing images. The model produced pixel-wise probability maps for each target region, and final masks were constructed based on sigmoid activation functions and thresholded to 0.5. Segmentation accuracy was evaluated with Dice and Jaccard overlap metrics comparing predicted masks against the provided testing masks. The model achieved a dice coefficient of 0.975 and a Jaccard index of 0.952 on the validation image set, with similar performance on the testing image set (Dice:0.968, Jaccard: 0.938). The slight drop in testing metrics indicates strong generalization with minimal overfitting. These metrics suggest reliable anatomical masking performance.

#### Image Analysis

Analysis was conducted utilizing a custom set of Python Scripts. Regional masks provided by our machine learning algorithm were used to restrict analysis to specific regions. Raw images were originally captured at 16 bit depth [intensity range:0-65535], and a high pass filter at 1500 was applied to cut out low intensity noise. After the high pass filter had been applied, puncta were identified and segmented to generate filtered images for further analysis. For each channel (CB1R, VGAT, VGLUT1, VGLUT2), puncta were segmented independently using a difference-of-Gaussians based detection pipeline to isolate noise, followed by watershed segmentation to define puncta centroids. Owing to the unique expression pattern of each protein, thresholds and parameters were adjusted for each protein, but were kept consistent across all genotypes and images. During parameter optimization, investigators were blinded to genotype.

After puncta had been processed, the following properties were computed for each channel: puncta area, intensity (mean, median, maximum), and puncta density (normalized to molecular layer area per image). Cross channel colocalization metrics were computed to examine interaction of proteins. Throughout the results, puncta colocalization was determined by counting synaptic puncta (VGAT, VGLUT1, or VGLUT2) with at least one CB1R puncta within 0.7 µm, and dividing that number by the total detected synaptic puncta to give a % colocalization metric. Mean intensity of both colocalized and non-colocalized puncta was also recorded. To validate that colocalization represented true coincidence between signals and not simply random overlap, we also calculated % colocalization between CB1R and the three vesicular markers after shifting the CB1R channel by 100 pixels along the X axis. Percent colocalization for all three synaptic markers was significantly reduced in the pixel shifted images (Supplemental Fig. 1D-F), confirming true colocalization of CB1R with each synaptic marker. The complete data-set of all measured values is available in the supplemental table.

Investigators were blinded to genotype throughout the analysis. Analysis code was written in Python 3.11, utilizing NumPy (BSD), SciPy (BSD), Pandas (BSD), scikit-image (BSD), tifffile (MIT), matplotlib (PSF), and tqdm (MPL 2.0) libraries. These open-source libraries were used with no restrictions on academic research and were cited according to publisher guidelines.

### Statistical Analysis

All data are presented as mean +/− SEM. Statistical significance and p-values were determined in GraphPad Prism using two-way ANOVAs and unpaired Welch’s t-tests. Reported n-values represent the number of cells or images analyzed. ns (p-value>0.05), * (p-value≤0.05), **(p-value≤0.01), ***(p-value≤0.001), ****(p-value≤0.0001).

## RESULTS

Given the importance of CB1R signaling in regulating neuronal circuits (Busquets-Garcia and Bains, 2018), the dysregulation of these receptors in peripheral tissue in DMD (Iannotti et al., 2018), and the normally high expression of both dystrophin and CB1R in cerebellum (Lidov et al., 1994; Kawamura et al., 2006), we first examined CB1R expression in the cerebellar cortex by immunofluorescent labeling in adult (P50-60) DMD^mdx^ mice and WT littermate controls (Fig. 1A).

**Figure 1:**
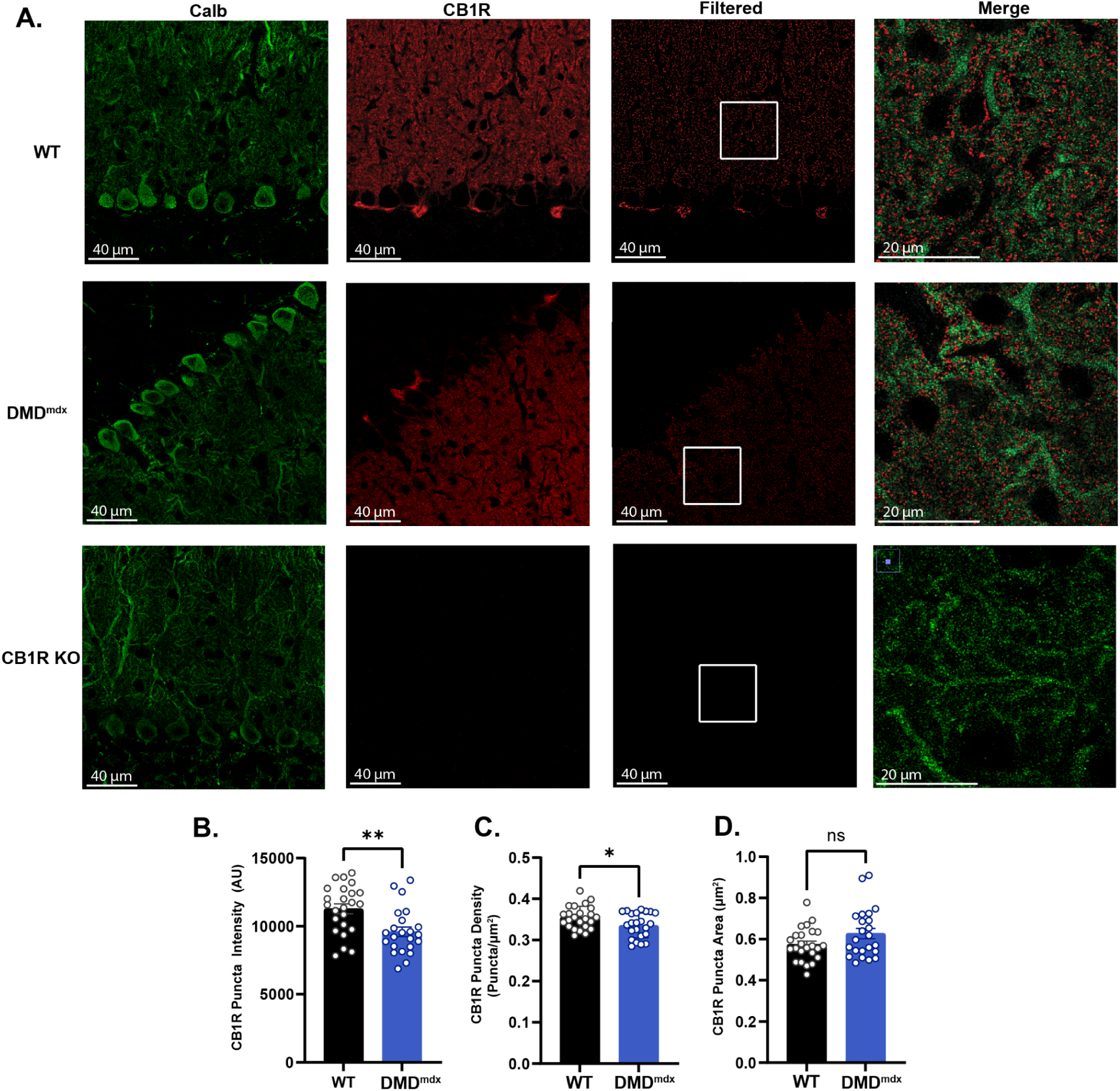
CB1R labeling is altered in DMD^mdx^ mice. **A.** Representative confocal images from WT, DMD^mdx^ and CB1R KO cerebella demonstrating calbindin (green), raw CB1R (red), and filtered CB1R puncta (see methods: image analysis). The right column shows an expanded and merged view of the area indicated in the filtered CB1R column. Quantification of mean CB1R puncta intensity (**B**), density (**C**), and area (**D**) in WT (black) and DMDmdx (blue) cerebella.

We found that CB1R puncta intensity in the molecular layer of the cerebellum is significantly reduced in DMD^mdx^ mice (WT: 11,282±359, n=24; DMD^mdx^: 9,596±352, n=23; *p*=0.0016; Fig. 1B), with a slight reduction in puncta density (WT: 0.35±0.005 puncta/μm^2^, DMD^mdx^: 0.335±0.006 puncta/μm^2^, *p*=0.028, Fig. 1C), demonstrating the endocannabinoid system is also altered in the CNS. Puncta area was not significantly different across genotypes (WT: 0.58±0.02 μm^2^, DMD^mdx^: 0.63±0.03 μm^2^, *p*=0.11, Fig. 1D). Within the cerebellum, CB1Rs are expressed at a variety of synapses, which may show differential expression and modulation of receptor populations. CB1Rs are most highly expressed at presynaptic terminals; in the molecular layer of the cerebellum, this includes expression at presynaptic boutons of MLIs, PF, and CFs (Kawamura et al., 2006). Each of these synapses play unique roles in regulating PC activity and synaptic plasticity. We therefore sought to determine how CB1R expression is modulated at each of these synapses independently due to loss of dystrophin expression.

### CB1R expression at MLI-PC synapses

Previous studies have shown that dystrophin is selectively expressed at the postsynaptic densities of inhibitory synapses between MLIs and PCs in the cerebellum (Knuesel et al., 1999; Briatore et al. 2020; Wu et al., 2022). These synapses also express high levels of CB1Rs (Diana et al., 2002; Kawamura et al., 2006), leading us to hypothesize that disruption of CB1R expression at these synapses may be responsible for the decrease in total CB1R expression observed in the molecular layer (Fig. 1B,C). To test this possibility, we co-labeled cerebellar sections for CB1R and VGAT, a marker of inhibitory synapses (Fig. 2A).

**Figure 2:**
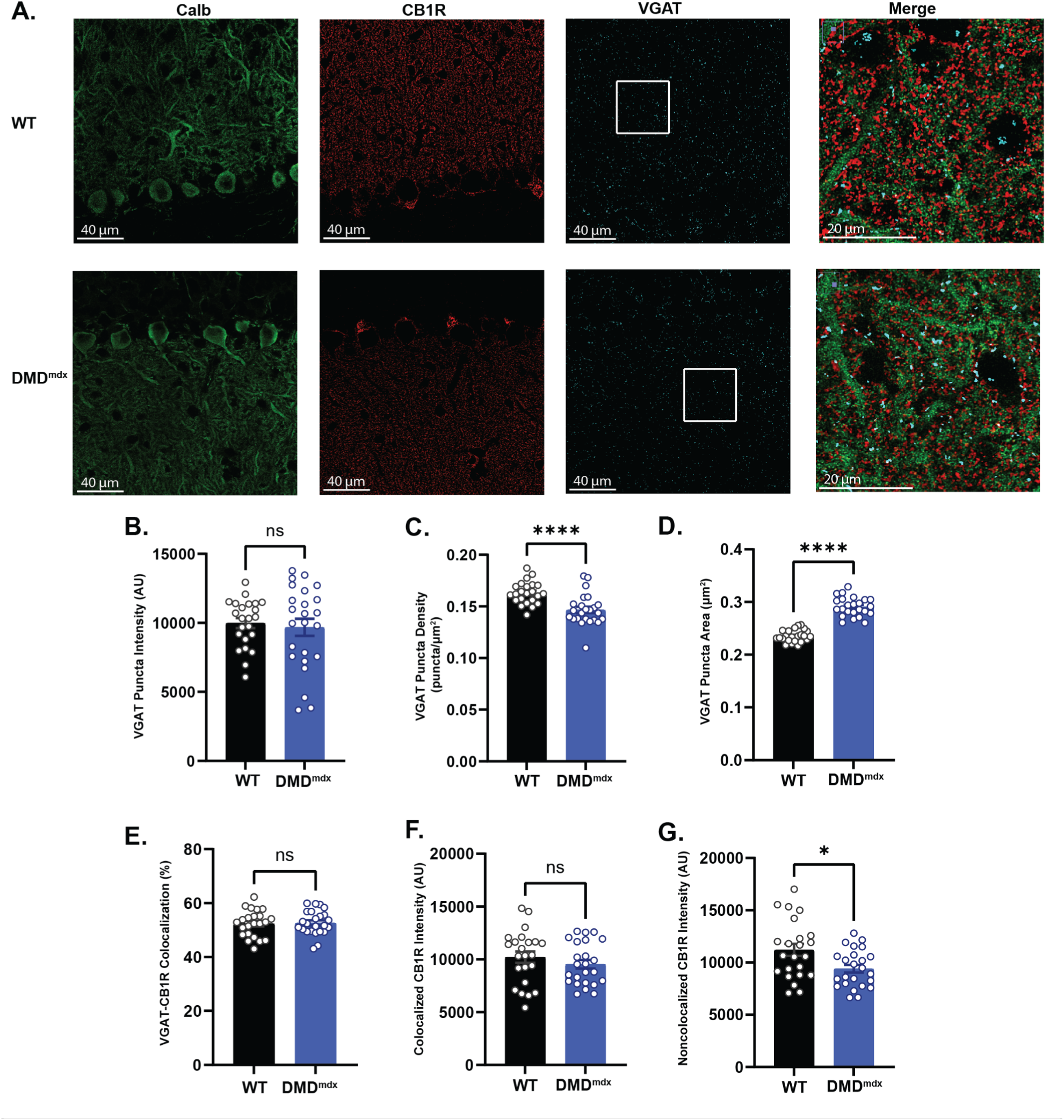
Colocalization of VGAT and CB1R. **A.** Representative images demonstrating calbindin (green), filtered CB1R (red), and filtered VGAT (cyan) staining across WT and DMD^mdx^ cerebella. The right column shows an expanded and merged view of the area indicated in the VGAT image. Quantification of mean intensity (**B**), density (**C**), and area (**D**) of VGAT puncta in the molecular layer. **E.** Percentage of VGAT puncta colocalized with CB1R puncta. **F.** Intensity of CB1R puncta colocalized with VGAT puncta. **G.** Intensity of CB1R puncta not colocalized with VGAT puncta.

First, we quantified properties of VGAT puncta in isolation - previous work has established that loss of dystrophin can impact synaptic inhibition and reduce the number of inhibitory synapses (Wu et al., 2023). Consistent with this literature, we find a ∼10% reduction in VGAT puncta density throughout the molecular layer of DMD^mdx^ mice compared to WT controls (WT: 0.163±0.002 puncta/μm^2^, n=23; DMD^mdx^: 0.147±0.003 puncta/μm^2^, n=24; *p*<0.0001, Fig. 2C). Curiously, we also observed a significant increase in the average size of VGAT puncta (WT: 0.236±0.002 μm^2^, n=23; DMD^mdx^: 0.291±0.004 μm^2^, n=24; *p*<0.0001, Fig 2D), though no difference in the average puncta intensity (WT: 9,978±360, n=23; DMD^mdx^: 9,676±615, n=24; *p*=0.68, Fig. 2B). This is surprising as previous work using electron microscopy has found decreased active zone length and vesicle number at MLI synapses onto the PC soma (Wu et al., 2022), possibly suggesting smaller synapses, though total bouton size was not measured. We did not observe a change in the intensity (WT: 9,036±424, n=23; DMD^mdx^: 8,294±588, n=24; *p*=0.31, data not shown) of VGAT labeling at pinceau synapses, another inhibitory synapse surrounding the axon initial segment of PCs. This is consistent with lack of dystrophin expression at pinceau synapses, and suggests that changes in VGAT labeling in the molecular layer are not simply due to differences in labeling efficiency across samples. In order to measure CB1R expression specifically at MLI boutons, we measured CB1R labeling intensity colocalized with VGAT+ puncta. We find no change in the percentage of CB1-VGAT pairs (VGAT+ puncta within 0.7 μm of a CB1R puncta, see methods) (WT: 52.5±1.03%, n=23; DMD^mdx^: 52.72±0.92%, n=24; *p*=0.84, Fig. 2E). Contrary to our initial hypothesis, we did not find a significant difference in colocalized CB1R intensity (WT: 10,210±531, n=23; DMD^mdx^: 9,535±408, n=24; *p*=0.32, Fig. 2F) across genotypes at inhibitory (VGAT+) synapses. These data suggest that while improper dystrophin expression reduces inhibitory synapse number, CB1R expression at remaining synapses is unaltered. However, we did see a significant decrease in CB1R intensity at CB1R puncta not colocalized with VGAT (WT: 11,202±584, n=23; DMD^mdx^: 9,411±369, n=24; *p*=0.014, Fig 2G), consistent with the overall decrease in labeling observed in the molecular layer (Fig. 1B), and suggesting CB1R expression is reduced at other synapses.

In order to assess the functional activity of CB1Rs at MLI-PC synapses, we measured depolarization-induced suppression of inhibition (DSI) in PCs of WT and DMD^mdx^ mice. DSI is a form of short-term plasticity mediated by CB1Rs (Diana et al., 2002; Kawamura et al., 2006), making it an effective measure of functional CB1R activation. We did not find a difference in the magnitude of DSI between genotypes (WT: 60.8±4.6% of baseline, *n*=12; DMD^mdx^: 59.3±5.6% of baseline, *n*=13; *p*=0.84, Fig. 3B-C), indicating CB1R activation at MLI synapses is similar. Taken together, these data demonstrate that despite the localization of dystrophin to inhibitory synapses and the substantial decrease in inhibitory synapse number in DMD^mdx^ PCs (Fig. 2C), endocannabinoid signaling and CB1R expression are unaltered at MLI synapse. This raises the question of what accounts for the loss of total CB1R expression seen in the molecular layer.

**Figure 3:**
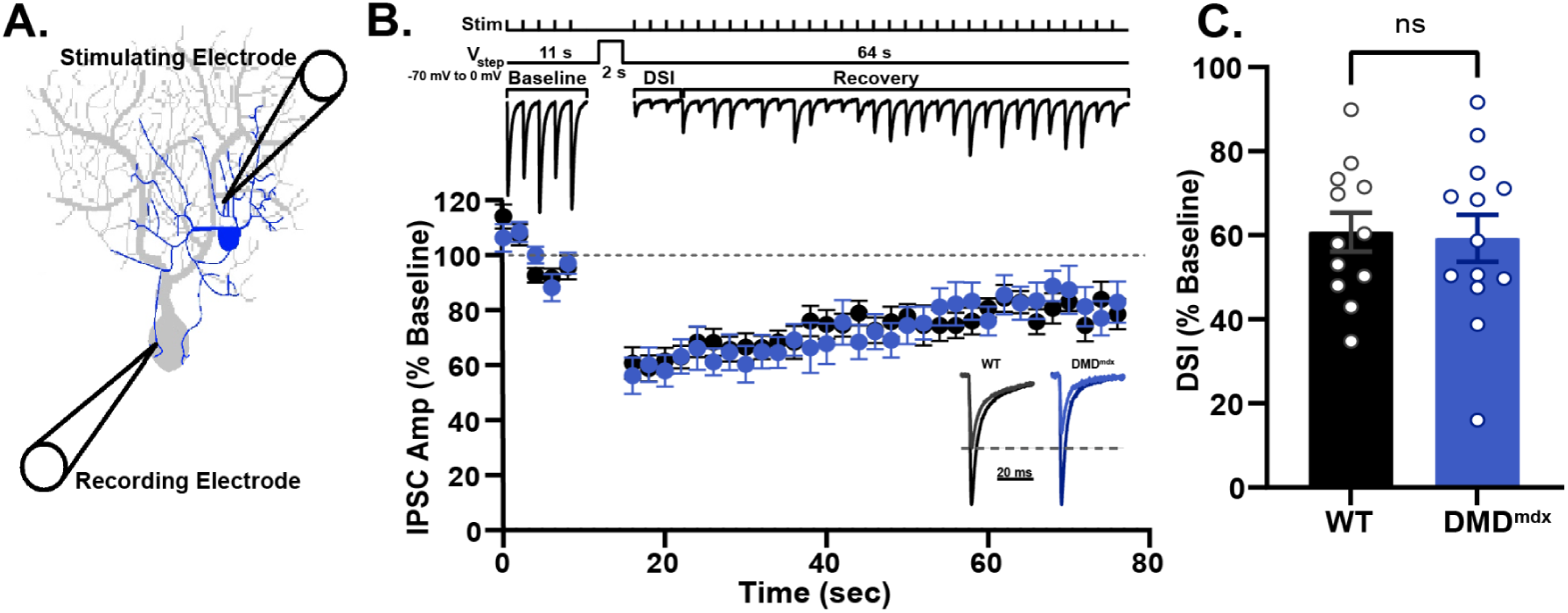
DSI is not significantly different at the MLI-PC synapse between WT and DMD^mdx^ mice. **A:** Diagram showing recording and stimulating electrode configuration for DSI experiments. **B:** *Top:* Diagram showing the timing of MLI stimulation (0.5 Hz) and PC depolarization (voltage step from −70 mV to 0 mV) relative to a representative trace of MLI-PC DSI. *Bottom:* Average IPSC amplitudes across the DSI protocol represented as percentage of baseline from WT (black) and DMD^mdx^ (blue) PCs. *Inset:* Representative traces of IPSCs at baseline (black/dark blue) and post-depolarization (grey/light blue) from WT (left) and DMD^mdx^ (right) PCs. **C:** Average magnitude of DSI represented as percentage of baseline in WT (black) and DMD^mdx^ (blue) PCs.

### CB1R expression at PF synapses

To address this question, we next examined PF synapses, the most numerous synapse type in the molecular layer. As in Figure 2, we labeled cerebellar sections for calbindin and CB1Rs, but in this case used the Vesicular Glutamate Transporter 1 (VGLUT1) antibody to specifically label PF synapses (Fig. 4A). We did not find a difference in VGLUT1 puncta intensity (WT: 5,053±258, *n*=24; DMD^mdx^: 4,805±218, *n*=23; *p*=0.47, Fig. 4B), density (WT: 0.437±0.006 puncta/μm^2^, *n*=24; DMD^mdx^: 0.425±0.004 puncta/μm^2^, *n*=23; *p*=0.10, Fig. 4C), or area (WT: 0.260±0.005 μm^2^, *n*=24; DMD^mdx^: 0.261±0.004 μm^2^, *n*=23; *p*=0.87, Fig. 4D), indicating that gross PF synapse number and morphology are not altered in DMD^mdx^ mice. However, we did find changes in CB1R expression at VGLUT1+ puncta. While the percentage of VGLUT1 puncta that colocalize with a CB1R puncta was not different (WT: 62.9±0.74%, *n*=24; DMD^mdx^: 61.8± 0.99%, *n*=23; *p*=0.42, Fig. 4E), we found that CB1R labeling intensity at VGLUT1 puncta is significantly reduced (WT: 11907±407, *n*=24; DMD^mdx^: 9,837±359, *n*=23; *p*=0.0004, Fig. 4F) in DMD^mdx^, suggesting a consistent downregulation of CB1Rs at PF synapses. The same samples showed no difference in CB1R labeling at pinceau synapses across genotypes (Intensity: WT: 18,904±693, *n*=24; DMD^mdx^: 18,091±454, *n*=23; *p*=0.33, Area: WT: 74.69±2.51 μm^2^, *n*=24; DMD^mdx^: 76.31±3.16 μm^2^, *n*=23; *p*=0.69; Supplemental Fig 1B,1C), suggesting CB1R labeling efficiency was similar across samples. Further, CB1R intensity at puncta not colocalized with VGLUT1 was not different between genotypes (WT: 9,748±306, *n*=24; DMD^mdx^: 9,023±332, *n*=23; *p*=0.12, Fig. 4G), consistent with the lack of change in CB1R labeling at VGAT puncta (Fig. 2F). This also provides further evidence that CB1R labeling efficiency was similar across genotypes.

**Figure 4:**
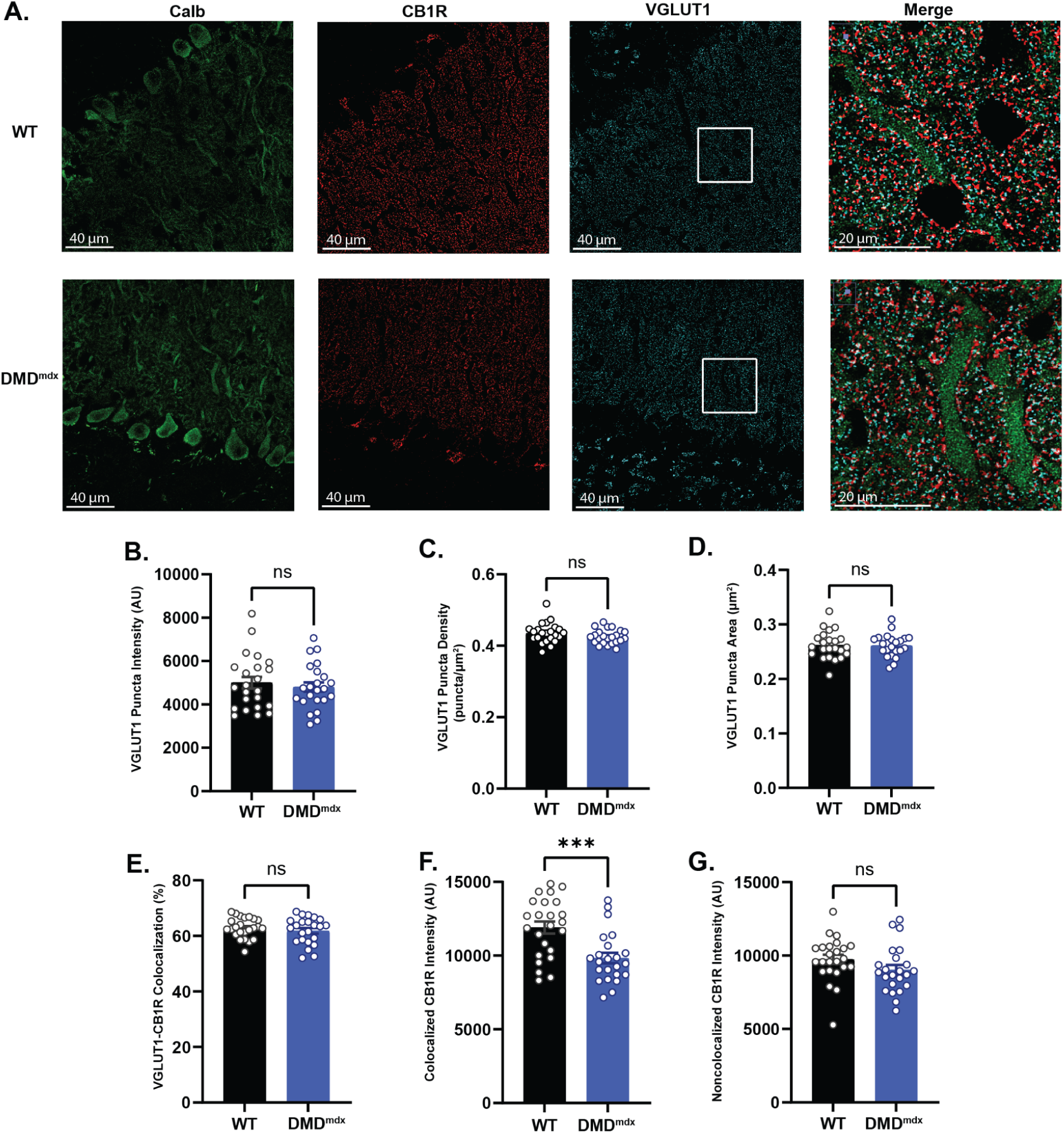
Colocalization of VGLUT1 and CB1R. **A.** Representative images demonstrating calbindin (green), filtered CB1R (red), and filtered VGLUT1 (cyan) staining across WT and DMD^mdx^ cerebella. The right column shows an expanded and merged view of the area indicated in the VGLUT1 image. Quantification of mean intensity (**B**), density (**C**), and area (**D**) of VGLUT1 puncta in the molecular layer. **E.** Percentage of VGLUT1 puncta associated with CB1R puncta. **F.** Intensity of CB1R puncta colocalized with VGLUT1 puncta. **G.** Intensity of CB1R puncta not colocalized with VGLUT1 puncta

In order to measure functional changes in endocannabinoid signaling at PF synapses, we measured DSE, a CB1R-dependent form of short-term plasticity similar to DSI (Kreitzer and Regehr, 2001). We found a significant reduction in the magnitude of suppression (WT:64.1±5.4 % of baseline, n=9; DMD^mdx^: 79.3±1.7 % of baseline, n=9, *p*=0.025; Fig. 5B-C) in DMD^mdx^ PCs, pointing towards functional deficits in CB1R-mediated signaling. There was no difference in paired-pulse ratio (PPR; WT: 1.46 ± 0.03, *n*=20; DMD^mdx^: 1.45 ± 0.02, *n*=19; *p*=0.73, Fig. 5D) or stimulus-response curves (for a two-way ANOVA, F=2.13 and p=0.16 between WT, *n*=18 and DMD^mdx^, *n*=13, Fig. 5E) of PF EPSCs across genotypes, suggesting the reduced DSE in DMD^mdx^ is not due to changes in baseline release probability. Interestingly, DSE in WT PCs was highly variable, ranging from ∼30 to 85% of baseline, while DSE in DMD^mdx^ was quite uniform, indicating that EC signalling may normally be highly heterogeneous, with some synapses displaying significant short-term plasticity and others very little. This heterogeneity is lost in the DMD^mdx^ context.

**Figure 5:**
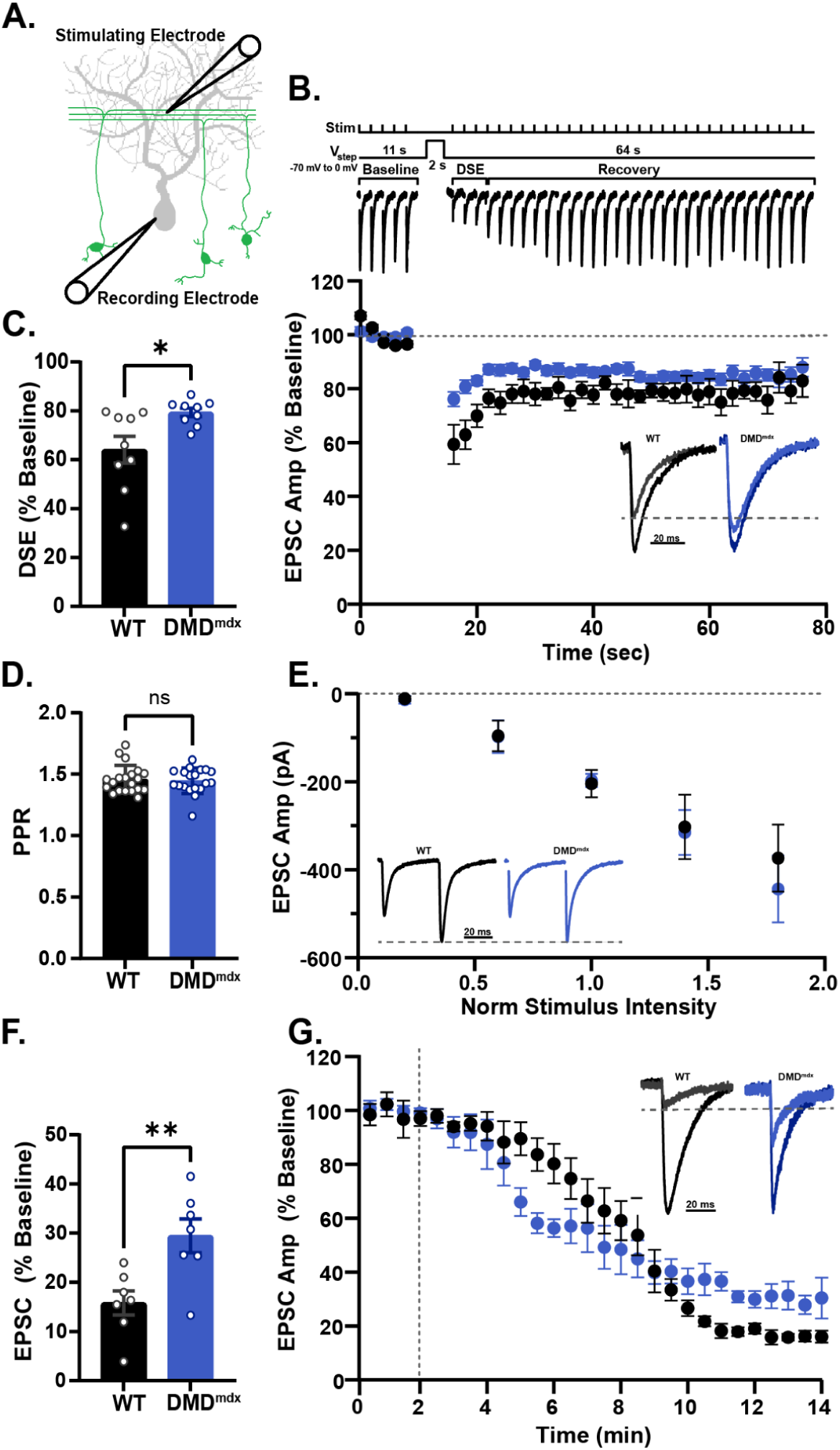
PF DSE is impaired in DMDmdx PCs. A. Diagram showing recording and stimulating electrode configuration for PF-DSE experiments. *B. Top:* Diagram showing the timing of PF stimulation (0.5 Hz) and PC depolarization (voltage step from −70 mV to 0 mV) relative to a representative trace of PF-PC DSE. *Bottom:* Average EPSC amplitudes showing DSE and recovery over time in WT (black) and DMD^mdx^ (blue). *Inset:* Representative traces of EPSCs at baseline (black/dark blue) and post-depolarization (grey/light blue) in WT (left) and DMD^mdx^ (right) PCs. **C.** Average magnitude of DSE represented as percent of baseline between WT (black) and DMD^mdx^ (blue) PCs. **D.** Average paired-pulse ratio (PPR) of PF-EPSCs from WT (black) and DMD^mdx^ (blue) PCs. **E.** Stimulus-response curves of PF-EPSC amplitudes over a range of stimulus intensities. *Inset:* Example traces of pairs of PF-EPSCs (50 ms ISI) from WT (black) and DMD^mdx^ (blue) PCs. **F.** Average change in PF-EPSC amplitude following WIN application in WT (black) and DMD^mdx^ (blue) PCs. **G.** Average EPSC amplitudes before and during WIN wash-on. Dotted line represents application of WIN to the bath. *Inset:* Example traces (normalized to the baseline amplitude) of PF-EPSCs before (black/blue) and after (grey/light blue) WIN application

Our DSE experiments point to a reduction in endocannabinoid signaling, but do not establish a mechanism of change, which could include reduced 2-AG release, reduced CB1R expression at PF boutons, or increased 2-AG removal from the synaptic space (Castillo et al., 2012). Our immunolabeling results (Fig. 4F) point toward a reduction in CB1R expression, and the lack of change in DSI at MLI-PC synapses (Fig. 3C) suggests that 2-AG production in PCs is not broadly altered. However, to isolate potential changes in CB1R expression/signaling, we eliminated 2-AG production and removal as variables and activated CB1R through bath application of WIN 55-212-2 (WIN, 1 μM), a CB1R agonist. We found that inhibition of PF EPSC amplitude in the presence of WIN was significantly reduced in DMD^mdx^ PCs (WT: 15.8±2.5% of baseline, n=7 DMD^mdx^: 29.5±3.5% of baseline, n= 7, *p*=.0083, Fig. 5F-G), suggesting reduced endocannabinoid signaling is primarily due to decreased CB1R expression at PF boutons.

Taken together, these data indicate that while the PF synapses are regularly formed and distributed with unchanged release properties, there appears to be a significant reduction in endocannabinoid signaling at the PF-PC synapse, consistent with reduced populations of functional CB1Rs.

### CB1R expression at CF synapses

CB1R expression has also been reported at climbing fiber synapses (Kawamura et al., 2006), a unique excitatory synapse arising from axon projections of the inferior olive. To identify possible changes in endocannabinoid signaling at this synapse, we labeled parasagittal slices for calbindin, CB1R, and VGLUT2, a vesicular marker specific to CF synapses (Fig. 6A). Quantification of VGLUT2 labeling revealed no change in puncta intensity (WT: 8,494±386, *n*=24; DMD^mdx^: 8,126±357, *n*=24; *p*=0.49, Fig. 6B), density (WT: 0.023±0.001 puncta/μm^2^, *n*=24; DMD^mdx^: 0.024±0.001 puncta/μm^2^, *n*=24; *p*=0.41, Fig. 6C), or area (WT: 0.567±0.014 μm^2^, *n*=24; DMD^mdx^: 0.581±0.14 μm^2^, *n*=24; *p*=0.46, Fig. 6D). CB1R labeling at VGLUT2 puncta was generally quite low compared to VGLUT1 or VGAT puncta, suggesting CB1R expression may be generally low at CF terminals. However, we did find slightly reduced CB1R intensity at VGLUT2 puncta in DMD^mdx^ compared to WT (WT: 8,570±544, *n*=24; DMD^mdx^: 7,034±377, *n*=24; *p*=0.025, Fig. 6F), though the proportion of VGLUT2+ puncta colocalizing with CB1R did not change (WT: 38.4±1.7%, *n*=24; DMD^mdx^: 36.8±1.2%, *n*=24; *p*=0.46, Fig. 6E). CB1R intensity at puncta not localized to VGLUT2 terminals was also reduced in DMD^mdx^ (WT: 10,157±633, *n*=24; DMD^mdx^: 8,326±400, *n*=24; *p*=0.019, Fig. 6G), consistent with our findings at PF (VGLUT1+) synapses (Fig. 4F). CB1R labeling in the pinceau region of VGLUT2 labelled sections was not different from labeling with other markers (VGAT and VGLUT1) and was not different across genotypes (Supplemental Fig. 1B), suggesting reduced CB1R intensity is not due to differences in labeling efficiency.

**Figure 6:**
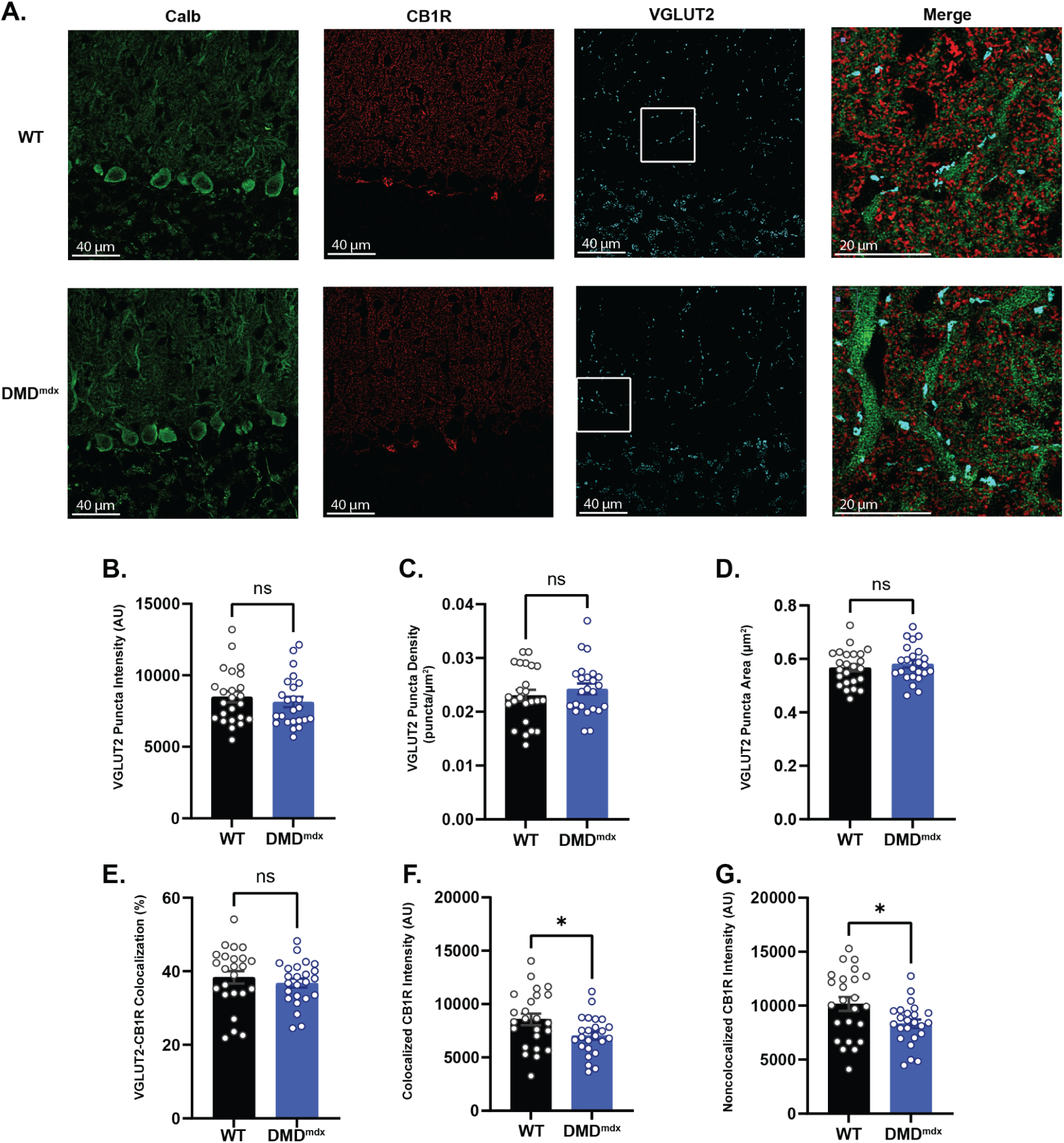
Colocalization of VGLUT2 and CB1R. **A.** Representative images demonstrating calbindin (green), filtered CB1R (red), and filtered VGLUT2 (cyan) staining across WT and DMD^mdx^ cerebella. The right column shows an expanded and merged view of the area indicated in the VGLUT2 image. Quantification of mean intensity (**B**), density (**C**), and area (**D**) of VGLUT2 puncta in the molecular layer. **E.** Percentage of VGLUT2 puncta associated with CB1R puncta. **F.** Average intensity of CB1R puncta colocalized with VGLUT2 puncta. **G.** Intensity of CB1R puncta not colocalized with VGLUT2 puncta.

In order to validate whether functional CB1R activity differs at CF synapses, we again measured CB1R-dependent short-term plasticity (Kreitzer and Regehr, 2001). In contrast with the reduced CB1R immunolabeling at CF boutons, we did not find a significant difference in DSE between WT and DMD^mdx^ (WT: 90.9 ± 1.9% of baseline, *n*=8; DMD^mdx^: 91.0 ± 2.4% of baseline, *n*=8; *p*=0.96, Fig. 7B-C). However, the magnitude of inhibition at CF synapses (∼90% of baseline) was quite small compared to either MLI-PC (∼60%) or PF-PC (∼65%) synapses (despite using a longer PC depolarization in CF DSE experiments, see methods), potentially obscuring subtle changes in CB1R activity at this synapse. To maximize CB1R activation at CF synapses and identify possible changes in CB1R expression, we measured inhibition of CF-EPSCs following bath application of WIN. In both genotypes, inhibition of CF-EPSCs following WIN application was small (∼70% of baseline) compared to similar experiments at PFs (Fig. 5F), consistent with the small magnitude of DSE observed at these synapses. Further, there was not a significant difference in the magnitude of inhibition from WIN across genotypes (WT: 68.7 ± 2.5% of baseline, *n*=7; DMD^mdx^: 67.7 ± 2.9% of baseline, *n*=7; *p*=0.79, Fig. 7D-E), suggesting there is no difference in CB1R activity at these synapses. Together, these data suggest that CB1R expression and activation are relatively weak at CF synapses, with no change in expression following the loss of dystrophin.

**Figure 7:**
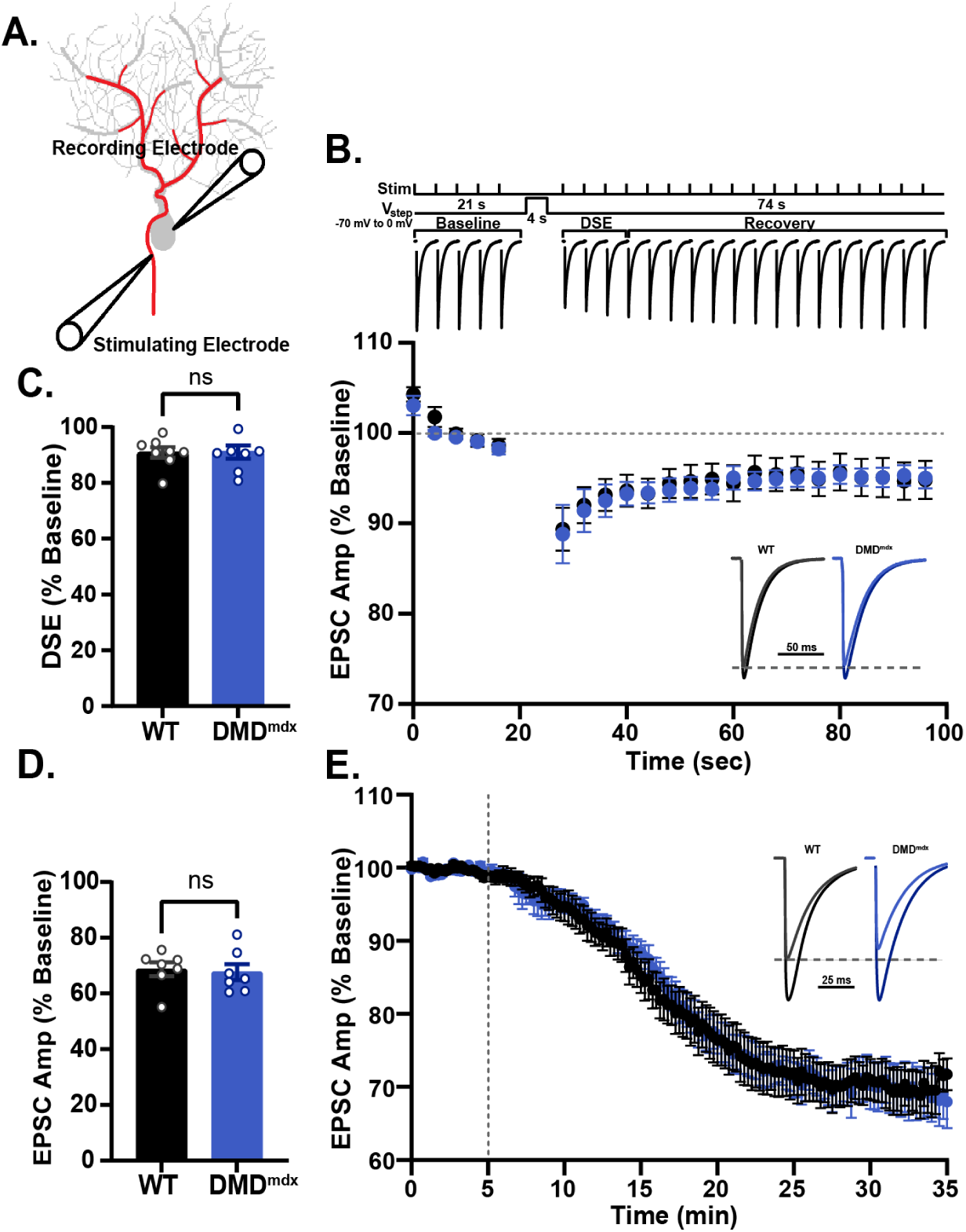
CF-DSE is not different in DMDmdx PCs. **A:** Diagram showing recording and stimulating electrode configuration for CF-DSE and WIN experiments. **B:** *Top:* Diagram showing the timing of CF stimulation (0.25 Hz) and PC depolarization (voltage step from −70 mV to 0 mV) relative to a representative trace of CF-PC DSE. *Bottom:* Average EPSC amplitudes showing DSE and recovery over time in WT (black) and DMDmdx (blue). *Inset:* Representative traces of EPSCs at baseline (black/dark blue) and post-depolarization (grey/light blue) between WT (left) and DMDmdx (right). **C:** Average magnitude of DSE represented as percentage of baseline between WT (black) and DMDmdx (blue). **D:** Average decrease in EPSC amplitude following WIN application as percentage of baseline between WT (black) and DMDmdx (blue). **E:** Average EPSC amplitude before and during bath application of 1 μm WIN represented as percentage of baseline in WT (black) and DMDmdx (blue). *Inset:* Representative traces of CF-EPSCs before (black/dark blue) and after (grey/light blue) WIN application.

### Long-term depression at PF synapses

Our data show a significant reduction in CB1R expression and functional activation at PF synapses in DMD^mdx^ mice. How does this change affect signaling and plasticity in the cerebellar circuit? Previous work has shown that CB1R activation is necessary for LTD induction at PF-PC synapses (Safo and Regehr, 2005; Carey et al., 2011) and cerebellar learning (Kishimoto and Kano, 2006; Steinmetz and Freeman, 2010). We therefore tested whether the decrease in CB1R expression at PF synapses is associated with a decrease in LTD induction. PF LTD was induced in WT and DMD^mdx^ PCs by repeated pairings of PF (10 stimuli at 100 Hz) and CF (2 stimuli at 20 Hz) stimulation (see methods, Fig. 8A). We found that LTD induction was robust in WT but completely absent in DMD^mdx^ PCs (WT: 71.4±6.5% of baseline, *n*=6;DMD^mdx^: 99.9±3.1% of baseline, *n*=7, *p*<0.005, Fig. 8B-C). These data suggest that reduced CB1R expression limits LTD induction at PF synapses. Though we cannot rule out the possibility that other changes caused by loss of dystrophin not explored here also contribute to loss of PF LTD, these data suggest that disruption of endocannabinoid signaling due to loss of dystrophin may play a significant role in the learning and cognitive deficits associated with dystrophin-deficiency.

**Figure 8:**
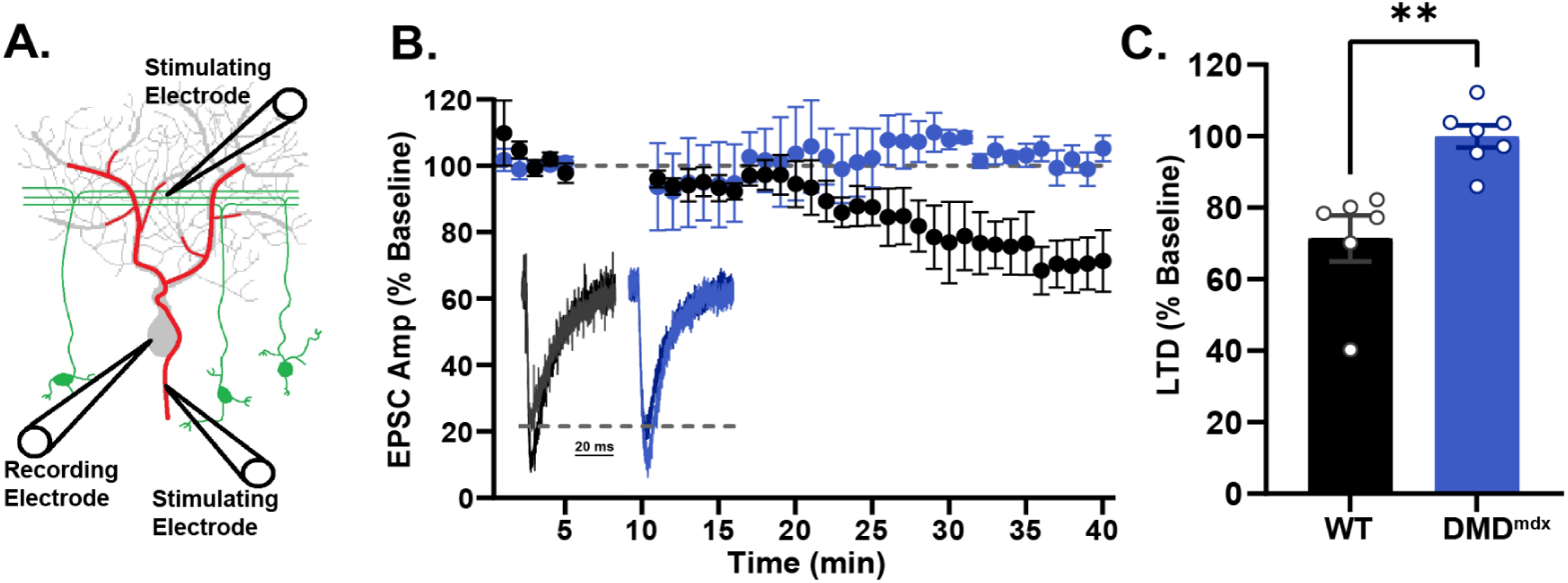
PF LTD is absent in DMDmdx PCs. **A:** Stimulus diagram showing electrode placement for LTD experiments. **B.** EPSCs over time showing baseline and post-induction magnitudes in WT (black) and DMDmdx (blue). *Inset:* Representative traces of EPSCs at baseline (black/dark blue) and post-LTD induction (grey/light blue) from WT (left) and DMDmdx (right) PCs. **C.** quantification of B, showing final magnitudes of depression in both genotypes.

## DISCUSSION

We find that CB1R expression and functional activation are reduced in the cerebellum of dystrophin-deficient DMD^mdx^ mice using a combination of immunofluorescent labeling and patch clamp electrophysiology. Our data using synapse specific markers show that CB1R expression is reduced at PF-PC synapses, but unchanged at MLI-PC or CF-PC synapses. Patch clamp electrophysiology experiments confirm functional downregulation of CB1Rs specifically at PF synapses. Further, we find that loss of CB1R expression reduces expression of long-term plasticity at PF synapses, potentially disrupting learning and adaptation in the cerebellar circuit. These data suggest changes in cannabinoid signaling may contribute to the cognitive and neurodevelopmental deficits associated with DMD.

In this study we measured CB1R expression at MLI, PF, and CF synapses using the synapse specific markers VGAT, VGLUT1, and VGLUT2, respectively. One potential caveat to this approach is that these markers do not distinguish between synapses made onto PCs and synapses made onto other cell-types in the molecular layer, primarily MLIs. Previous reports find that approximately 90-95% of PF synapses are formed onto PCs (Palay and Chan-Palay, 1974; Pichitornchai et al., 1994), suggesting that measuring VGLUT1 puncta is likely a reasonable representation of PF-PC synapses. On the other hand, inhibitory synapses formed onto MLIs may constitute up to ∼30-40% of all inhibitory synapses in the molecular layer (Lackey et al., 2024; Briatore et al., 2010), suggesting VGAT puncta are a mix of synapses onto PCs and MLIs. It is therefore possible that subtle changes in CB1R expression at MLI-PC synapses measured by immunolabeling may be masked by lack of change or opposing changes at MLI-MLI or PC-MLI synapses. However, our electrophysiology data, showing no change in DSI in DMD^mdx^ PCs, suggests there is no functional change in CB1R activity. At VGLUT2 puncta, marking CF synapses, we also observed a decrease in CB1R expression. However, CB1R labelling was quite low at VGLUT2 puncta compared to VGLUT1 or VGAT puncta. Given the low intensity of CB1R labeling at VGLUT2 puncta and the fact that only a small percentage of CB1R puncta colocalize with VGLUT2 puncta (Supplemental Fig. 1F), it is possible that much of CB1R labeling observed to colocalize with VGLUT puncta is in fact localized at the much more numerous PF (VGLUT1+) synapses. In this case, the decrease in CB1R labeling observed at VGLUT2+ puncta likely reflects loss of CB1Rs at PFs, not a true decrease at CF synapses. In fact, our immunofluorescent labeling and electrophysiological data suggest that CB1R expression is quite low at CF synapses. These data, together with our electrophysiological data, point to significant changes in CB1R expression at PF synapses, but not MLI or CF synapses onto PCs.

There is increasing evidence that dynamic regulation of CB1R expression plays an important role in both normal and pathophysiological conditions. Changes in CB1R expression have been observed in several pathophysiological conditions. CB1R expression is increased in the hippocampus in rodent models of epilepsy (Karlócai et al., 2011; Bojnik et al., 2012) and in patients with temporal lobe epilepsy (Goffin et al., 2011). Likewise, CB1R expression is increased in early stages of Alzheimer’s disease, but then decreases at later stages (Manuel et al., 2014; Talarico et al., 2019). Finally, post-mortem studies of patients with spinocerebellar ataxia type 3 (SCA-3) and mouse models of SCA-3 have revealed a significant increase in CB1R expression in the cerebellum (Rodriguez-Cueto et al., 2014, 2016). It is currently unclear whether these changes play a pathogenic role or represent compensatory changes in response to circuit dysfunction elsewhere. In addition to pathophysiological changes in CB1R expression, recent evidence suggests CB1R expression is dynamically regulated during normal synapse and circuit activity. Previous studies have shown rapid, activity-dependent down-regulation of CB1Rs at presynaptic terminals following agonist application (Mikasova et al., 2008), 4 Hz PF stimulation (Yang et al., 2019), or PF LTD (Hunley et al., 2023), suggesting CB1R expression may be a plastic property of the synapse. The current work extends dysregulation of CB1R expression into a new range of diseases characterized by loss of dystrophin and associated proteins, and strengthens the view that plasticity of CB1R expression is a critical aspect of synapse function.

In cerebellar PCs dystrophin expression is limited to the postsynaptic densities of inhibitory synapses on the soma and dendrites (Knuesel et al., 1999; Briatore et al., 2020). Due to this distribution, studies have generally focused on changes in inhibition following loss of dystrophin. We and others have shown that loss of dystrophin significantly reduces the number, active zone length, vesicle number, and GABA_A_R clustering at inhibitory synapses (Knuesel et al., 1999; Kueh et al., 2011; Wu et al., 2022). This loss of phasic inhibition is associated with an increase in tonic inhibition mediated by extrasynaptic δ-subunit containing GABA_A_Rs (Mitra and Pugh, 2026). In contrast, few studies have reported changes at excitatory synapses. In fact, in the present study we find no change in the intensity and number of VGLUT1+ puncta (Fig. 4B-C) or the baseline release probability and stimulus-response curve of PF synapses (Fig. 5D-E). It is therefore surprising to find that changes in CB1R expression are exclusive to PF-PC synapses. What is the mechanistic link between loss of dystrophin at inhibitory synapses, reduced inhibitory synapses function, and altered PF endocannabinoid signaling and synaptic plasticity? Previous studies have shown that PF plasticity is highly sensitive to the level of inhibition in PCs. Paired PF-CF activity in the absence of inhibition results in robust LTD induction, while the same pairing in the presence of inhibition does not show LTD, and can even produce LTP (Rowan et al., 2018). This raises the possibility that the severely reduced inhibition observed in DMD^mdx^ PCs causes excessive LTD at PF synapses, potentially saturating the LTD mechanism (Lev-Ram et al., 2003; Nguyen-Vu et al., 2017; Shakhawat et al., 2024). Our recent work shows a down-regulation of CB1R expression at PF synapses following 4 Hz stimulation or following LTD induction (Yang et al., 2019; Hunley et al. 2023). We propose a model where loss of dystrophin in DMD^mdx^ PCs causes the loss and/or weakening of inhibitory MLI synapses onto PCs, resulting in exaggerated or extraneous LTD induction at PF synapses. Excessive LTD at PF synapses then produces a significant down-regulation of CB1R expression at PF synapses and saturation of LTD mechanisms. This model is consistent with our observations of reduced CB1R expression and reduced/absent LTD at PF synapses in DMD^mdx^ PCs. However, we cannot rule out other possible mechanisms connecting loss of dystrophin to PF CB1R expression, including increased motor error signaling in the cerebellar network, changes in input pathways due to loss of dystrophin in other brain regions, or loss of dystrophin from non-neuronal cells (Aranmolate et al., 2017). Further experiments including cell type specific dystrophin mutants and manipulation of inhibitory synapse function will be required to fully understand the mechanisms producing down-regulation of CB1R expression at PF synapses.

DMD is associated with a range of cognitive and neurodevelopmental deficits (Cotton et al., 2005). While CNS-related aspects of this disease cause significant burdens for patients and families, there is a lack of approved or proposed therapies targeting the CNS. Modulation of the endocannabinoid system has shown promise in treating muscular aspects of the disease, mainly through promoting myotube formation and reducing inflammation (Iannotti et al., 2019; Argenziano et al., 2023; Ferreira et al., 2025). However, use of cannabidiol or other cannabinoid compounds to treat CNS-related aspects of the disease have not been investigated in detail. Our data show that endocannabinoid signaling in the CNS is disrupted in DMD, potentially leading to loss of short- and long-term synaptic plasticity and altered circuit dynamics. This work describes a previously unappreciated aspect of CNS dysfunction in DMD, and raises the possibility that modulation of the cannabinoid system may alleviate or treat CNS-related aspects of this disease. Further investigations modulating cannabinoid signaling in the cerebellar circuit will be necessary to validate CB1Rs or other cannabinoid proteins as potential therapeutic targets.

## Acknowledgement

We are grateful to members of the Pugh lab for helpful discussion and comments on data collection and analysis. This work was funded by National Institutes of Health, Grant # R01 NS123933 and R01NS134868.

---

**Supplement 1:**
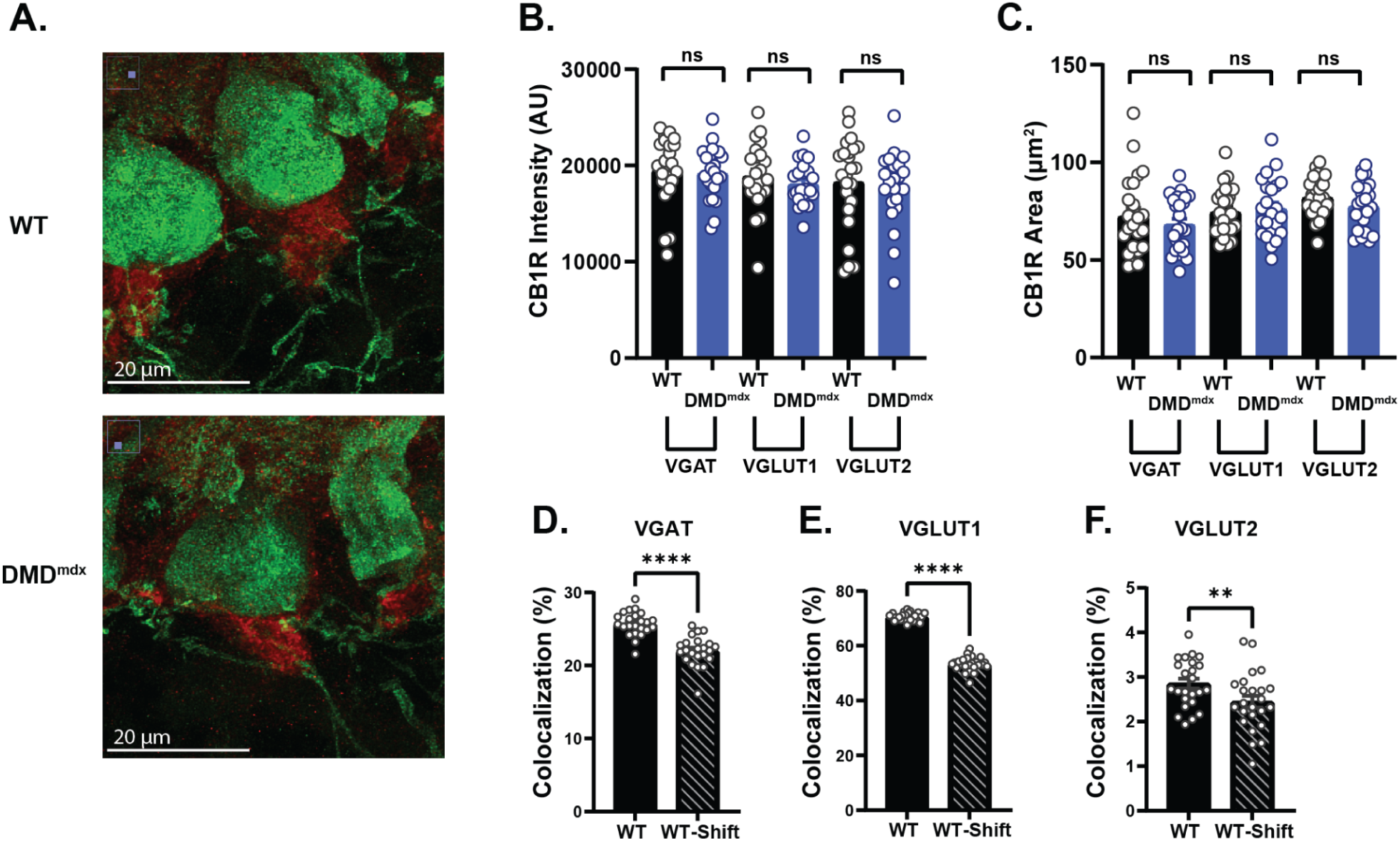
Validation of CB1R immunofluorescent labeling. **A.** Representative Pinceaux staining showing a PC Soma (green, calbindin) along with the characteristic pinceaux formation on the axon side (red, CB1R) in both genotypes. **B.** Quantification of average CB1R pinceaux intensity across both genotypes and slide sets (VGAT, VGLUT1, VGLUT2). **C.** Same as B, but area of pinceaux. **D-F.** Proportion of CB1R puncta that colocalize with the respective vesicular marker before and after a 100 pixel shift of the CB1R channel along the X axis.

